# Quilting the Brain: Whole-Brain iEEG Reconstruction via Incomplete Observation Linear Mixed Models

**DOI:** 10.64898/2026.05.31.729074

**Authors:** Ying Wang, Min Li, Maria L. Bringas-Vega, Pedro A. Valdes-Sosa

## Abstract

Mapping human brain function at high spatiotemporal resolution is constrained by the physical limitations of non-invasive imaging and the sparse sampling of invasive electrophysiology. While intracranial electroencephalography (iEEG) captures local eld potentials with millimeter precision, clinical implantation strategies result in a “coverage paradox” : observations are restricted to disjoint, patient-specific patches, leaving most of the cortex unobserved. This study introduces the Incomplete Observation Linear Mixed-Effect Model (IOLMM), a statistical framework that resolves this paradox by “quilting” fragmented observations into continuous, whole-brain source activity maps. Our approach integrates two innovations: (1) Sure Independence Screening (SIS) adapted from ultra-high-dimensional statistics to distinguish true physiological signals from volume-conducted “ghost sources”; (2) a hierarchical IOLMM that decouples group-level physiological fixed effects from subject-specific instrumental random effects, solving the scaling ambiguities that plague iEEG group analyses. Applied to the MNI Open iEEG Atlas, the framework is validated through sleep stage-dependent cortical source power reconstruction across Wake, N2, N3, and REM states, recovering the frontal predominance of NREM slow-wave activity and the graded electrophysiological hierarchy from fragmented recordings of 106 patients. This work establishes the first cortical surface-level normative electrophysiological atlas derived from iEEG, providing a quantitative reference for detecting and predicting epileptogenic lesions and bridging the gap between the microscopic precision of electrophysiology and the macroscopic scope of systems neuroscience.

**Highlights:**

- **Solving the Coverage Paradox:** A novel IOLMM statistical framework quilts sparse, non-overlapping iEEG data into a unified whole-brain probabilistic map.
- **Geometric Screening:** Adapts high-dimensional Sure Independence Screening (SIS) using cortical geometric eigenmodes to effectively filter out spurious ghost sources in inverse solutions.
- **Robust Harmonization:** Decouples group-level physiological fixed effects from subject-specific random effects, resolving scaling ambiguities inherent in heterogeneous multi-center recordings.
- **Biological Validation:** Validates the framework on real multi-center iEEG data by reconstructing sleep stage-dependent cortical source power maps, recovering known electrophysiological signatures of NREM and REM sleep from fragmented recordings.

## 1 Introduction

### 1.1 The Coverage Paradox in Brain Functional Mapping and the Renaissance of Intracranial Electrophysiology

The grand objective of Human Brain Mapping is to parse the complex spatiotemporal dynamics of neuronal activity, thereby elucidating the neural mechanisms underlying cognition, perception, and pathological states. Historically, this field has been dominated by non-invasive imaging technologies such as functional Magnetic Resonance Imaging (fMRI), Magnetoencephalography (MEG), and scalp Electroencephalography (EEG) (He et al., 2019; Friston, 2011). These technologies offer whole-brain coverage, enabling researchers to infer large-scale functional network architectures and global dynamic propagation (Sanz-Leon et al., 2015; Breakspear, 2017; Deco et al., 2008). However, these modalities face insurmountable biophysical limitations: scalp EEG and MEG suffer severe attenuation due to the volume conduction effect, where the skull and scalp act as low-pass filters, resulting in the loss of high-frequency signals and blurred source localization (Haufe et al., 2013; Van de Steen et al., 2019; Gonzalez-Moreira et al., 2020); conversely, fMRI relies on the hemodynamic response, the temporal resolution of which lags behind neuronal firing by seconds, failing to capture millisecond-scale neural dynamics (Bailes et al., 2023).

In contrast, Intracranial Electroencephalography (iEEG), including Stereoelectroen-cephalography (SEEG) and Electrocorticography (ECoG), offers an unparalleled window for observing the human brain with clarity. By implanting electrodes directly into deep brain tissue or placing them on the cortical surface, iEEG bypasses the filtering effects of the skull, capturing local field potentials (LFP) with millimeter-level spatial precision and millisecond-level temporal resolution, including high-frequency oscillations (HFOs) that provide direct access to local neuronal population dynamics (Parvizi and Kastner, 2018; Mercier et al., 2022; van Blooijs et al., 2023). This makes iEEG the “gold standard” for validating non-invasive findings, widely used to reveal fine spectral characteristics of visual processing streams (Vidal et al., 2010; Hamamé et al., 2014; Moraresku et al., 2025) and the propagation mechanisms of epileptic discharges (Smith et al., 2016; Sinha et al., 2017).

However, this high-fidelity signal comes with an inherent limitation—the “Coverage Paradox”. Unlike the uniform sensor arrays of MEG or the regular voxel grids of fMRI, iEEG sampling strategies are dictated entirely by clinical needs, typically used for presurgical evaluation in drug-resistant epilepsy patients to define the seizure onset zone (SOZ) and functional cortex (Parvizi and Kastner, 2018; Mercier et al., 2022). This clinically oriented implantation strategy results in a “spotlight” mode of observation: while it illuminates specific cortical patches, the vast majority of the brain remains in darkness (Parvizi and Kastner, 2018). This inherent sparsity means that no single subject can provide a complete map of whole-brain activity.

Consequently, the core challenge in iEEG research is how to aggregate these fragmented, non-overlapping, and spatially discrete individual observations into a coherent group-level whole-brain physiological atlas. Prior efforts have addressed this through channel-level normative mapping (Frauscher et al., 2018) or region-of-interest aggregation (Taylor et al., 2022; Pidnebesna et al., 2022), but these approaches operate in sensor space and sacrifice the spatial resolution that makes iEEG valuable. In statistics and signal processing, this aggregation challenge is often referred to as the “quilting problem” (Vinci et al., 2023; Chang et al., 2023). This is not merely a data stitching issue but a complex “Incomplete Observation Inference” problem, as the data missingness is structural rather than random. Solving this requires transcending traditional channel-level averaging to establish statistical models capable of handling structural missingness and extreme heterogeneity. This paper aims to propose a framework based on the Incomplete Observation Linear Mixed-Effect Model (IOLMM), which reconstructs a whole-brain neural activity map by “quilting” sparse source-space estimates onto a unified group manifold.

In constructing an iEEG group atlas, researchers must overcome two distinct types of heterogeneity: spatial sampling heterogeneity and signal amplitude heterogeneity.

### 1.2 Spatial Inconsistency and the Ill-Posed Inverse Problem

In traditional neuroimaging group analysis, spatial normalization is typically used to register individual brains to a standard stereotactic space (e.g., MNI152), enabling voxel-level cross-subject comparison (Mercier et al., 2022; Frauscher et al., 2018). However, for iEEG, even after spatial registration, electrode distribution between patients exhibits immense randomness and variance (Pidnebesna et al., 2022). One patient may have dense depth electrode coverage in the medial temporal lobe and orbitofrontal cortex, while another may only have electrodes in the operculum and insula. This spatial discrepancy renders traditional region of interest (ROI)-based averaging methods suboptimal. While computationally convenient, such approaches inherently necessitate a sacrifice of spatial resolution, thereby blurring functional boundaries and neglecting the fine-grained geometric orientation of neural generators (Taylor et al., 2022).

To preserve spatial resolution and bridge coverage gaps, recent research has turned to Electrophysiological Source Imaging (ESI) techniques (Cho et al., 2011) adapted for iEEG. ESI attempts to infer current density on the cortical surface from limited sensor voltage measurements by solving the inverse problem. However, the iEEG inverse problem possesses unique ill-posedness. Unlike scalp EEG sensors that surround the volume conductor, iEEG sensors are embedded within it, making them extremely sensitive to near-field sources while remaining virtually “blind” to far-field sources. Although algorithms like exact low-resolution brain electromagnetic tomography (eLORETA) are widely used for their zero-error localization properties, under sparse and irregular electrode arrays, the lead eld matrix—the physical model quantifying each source’s contribution to each sensor—becomes highly sensitive to individual implantation schemes. Without rigorous spatial screening, inverse solutions often yield numerous “ghost sources”^1^, which are spurious activations caused by noise or volume conduction far from the actual recording sites.

### 1.3 Signal Amplitude Scaling Discrepancies and the Failure of Global Normalization

Beyond spatial fragmentation, signal amplitude normalization presents a more insidious but fatal challenge. In scalp EEG analysis, applying a global scale factor (GSF) to normalize spectral energy across subjects is standard practice (Hernandez et al., 1994). The theoretical basis for GSF assumes that under a standardized sensor array (whole-scalp coverage), the total electric field energy recorded across individuals is roughly equivalent, with differences stemming primarily from non-physiological factors like skull thickness, skin impedance, or amplifier gain.

However, this assumption breaks down in the context of iEEG. Because the number of electrodes and the brain volume they cover are random and heterogeneous for each patient, the recorded “total energy” is effectively a function of sampling density, not the physiological state of the brain. A patient implanted with 150 electrodes covering a broad epileptic network will naturally record substantially higher total power than a patient with only 50 electrodes implanted to probe focal cortical dysplasia. Forcing a GSF normalization based on observed channel means in this scenario introduces systematic bias: more densely sampled regions or individuals may be “compressed” to a greater extent, causing true physiological strengths to be obscured by sampling bias.

Furthermore, the physical properties of the electrodes themselves (e.g., contact surface area, impedance) and their distance from neural generators (located in gray matter vs. white matter) introduce subject-specific or even channel-specific multiplicative scaling factors. Existing normalization methods, such as calculating percent signal change relative to a pre-stimulus baseline or Z-scores, are effective in event-related designs but are insufficient for resting-state analysis or atlas construction, due to the lack of a unified reference baseline for comparing “absolute” activation levels.

### 1.4 Statistical Framework for Incomplete Data: From Imputation to Mixed-Effects Inference

From a statistical perspective, the problem of constructing a whole-brain atlas from sparse observations shares theoretical connections with “matrix completion” or “graph quilting” (Chang et al., 2023; Vinci et al., 2023). In these frameworks, the goal is to infer missing entries in a covariance matrix or data tensor using observed local correlation structures. It is important to note that graph quilting targets *second-order* statistics (covariance/precision matrix estimation from block-missing data), whereas the present work addresses *first-order* statistics (estimating the population mean activation map from partially observed subjects). While low-rank matrix completion techniques show promise in calcium imaging and functional connectomics (Chang et al., 2023), they typically assume the data possesses a low-rank structure or stationarity, which may not hold for the high-dimensional and highly dynamic human brain source space. Moreover, pure imputation methods often ignore the hierarchical structure of the data—namely, that electrodes are nested within subjects—and the confounding of biological and technical sources of variation.

Linear Mixed Models (LMM) offer a more flexible and statistically rigorous alternative (Stroup, 2016). LMMs are widely used in neuroimaging to handle longitudinal data, repeated measures, and multi-level grouped designs. Their core advantage lies in the ability to decompose the variance of observed data into fixed effects and random effects. In the context of sparse iEEG, LMM allows us to model the observed source power at any cortical vertex as a linear combination of: true group-level physiological activity (fixed effects), subject-specific scaling biases (random effects), and measurement error.

Crucially, LMM does not require a balanced design; as long as the missing mechanism is reasonably modeled or adheres to the missing at random (MAR) assumption, LMM can provide unbiased parameter estimates even in the presence of substantial missing data. However, applying LMM to an iEEG source space with tens of thousands of vertices, while facing structural missingness (i.e., some subjects have absolutely no data in certain brain regions), requires specific model adaptation. This leads to the concept of the “Incomplete Observation Linear Mixed-Effect Model” (IOLMM), which introduces a binary selection matrix to explicitly map the sparse individual observation space to the dense group source space manifold.

### 1.5 High-Dimensional Screening and Geometric Eigenmode Constraints

Before utilizing IOLMM for “quilting”, the reliability of the input data must be ensured. As previously noted, the iEEG inverse problem is underdetermined, meaning infinite source configurations can explain the same set of sensor data. This uncertainty is amplified under sparse sampling, making the generation of “Ghost Sources” highly probable. If all reconstructed sources are directly input into the model, the group atlas will be overwhelmed by noise.

To address this, we must introduce screening techniques from high-dimensional statistics. Sure Independence Screening (SIS) theory, proposed by Fan and Lv (2008), proves that in ultra-high-dimensional regression problems (where the number of predictors *p* is far greater than the sample size *n*), screening based on marginal correlations can retain all important variables with probability approaching 1. In this study, we transfer this concept to source selection in ESI: by simulating smooth source activity that conforms to brain geometric structures and calculating the correlation between simulated sources and inverse solutions, we screen for “credible” source locations.

To generate biologically plausible simulated sources, we introduce cortical Geometric Eigenmodes. Recent research indicates that brain functional activity is strongly constrained by its anatomical geometry (Pang et al., 2023); by solving for the eigenfunctions of the cortical mesh’s Laplace-Beltrami Operator (LBO), one can obtain basis functions describing the brain’s intrinsic standing wave modes. Null models constructed based on these eigenmodes not only ensure the spatial continuity of simulated signals but also serve as a benchmark for screening the quality of real iEEG source reconstruction, thereby identifying “confident patches” where electrode coverage is sufficient and the solution is stable.

### 1.6 Research Objectives and Contributions

Addressing the “Quilting Problem” of multi-center iEEG data, this paper proposes a unified computational framework fusing Electrophysiological Source Imaging (ESI), high-dimensional variable screening, and Incomplete Observation Linear Mixed Models (IOLMM). Our method overcomes the limitations of traditional ROI averaging and interpolation by explicitly modeling the group mean and individual scaling factors in the signal generation process, allowing high-resolution whole-brain source activity reconstruction from structurally incomplete observations.

Specifically, the main contributions of this study are as follows:

1. Validating Screening Protocols: Adapting high-dimensional regression screening methods based on geometric eigenmode simulations (Fig. 1A) to identify reliable cortical vertices for source reconstruction under sparse sampling conditions (Fig. 1B), effectively eliminating ghost sources.
2. Establishing the IOLMM Framework: Constructing a linear mixed model containing subject-specific random effects (*b*_*i*_, to correct amplitude differences) and group-level fixed effects (***β***, representing the population-mean source power map), and solving parameters using the Expectation-Maximization (EM) algorithm to obtain the group-level activity of the whole brain instead of naïve average (Fig. 1C).
3. Evaluating Normalization Performance: Comparing the performance of scaling factors inferred by IOLMM against the traditional Global Scale Factor (GSF) under heterogeneous coverage, demonstrating the greater stability of the mixed-model approach in handling structurally missing data (Fig. 1D).
4. Biological Validation via Sleep Stages: Applying this pipeline to a large multi-center iEEG dataset from the Montreal Neurological Institute (MNI) (Frauscher et al., 2018), reconstructing sleep stage-dependent (Wake, N2, N3, REM) cortical source power maps from 106 patients with fragmented coverage, and recovering well-established electrophysiological signatures including the frontal predominance of NREM slow-wave activity (Fig. 1D).

**Figure 1:**
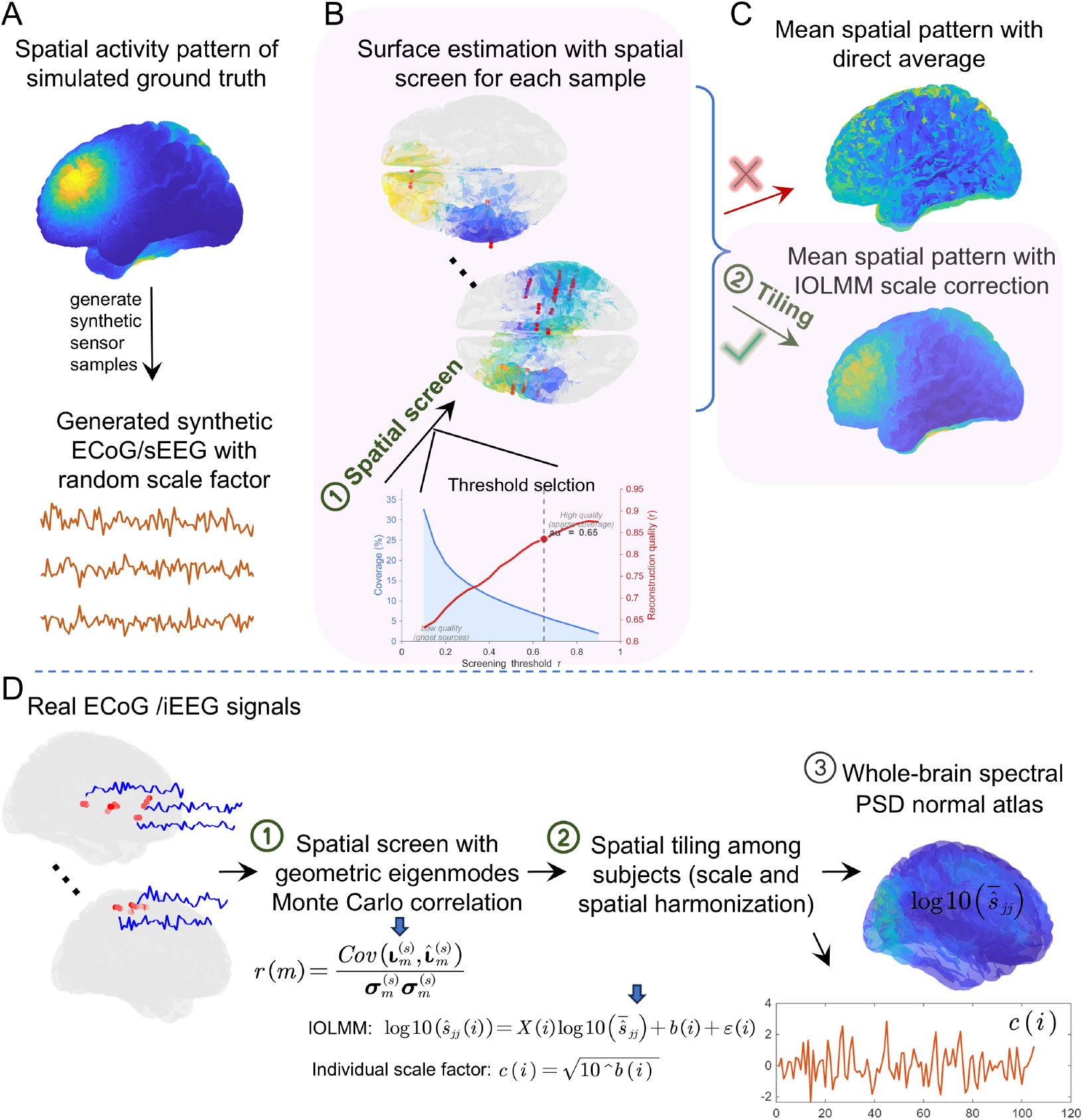
The whole diagram of brain quilting. A: Simulated ground truth mean surface activity and synthetic sensor samples randomly sampled from the forward model of the ground truth; B: Estimated source activity samples with spatial screening and the spatial preservation threshold selection; C: Comparison between simple sample average and quilting with IOLMM; D: Application to real iEEG data for individual scale estimation and sleep stage-dependent source power reconstruction.

This framework not only provides a solid mathematical foundation for group-level inference in iEEG data but also bridges the gap between micro-scale neuronal activity and macro-scale brain imaging atlases, with the potential for constructing normative electrophysiological references and localizing pathological anomalies.

## 2 Methods

Before formalizing the model, we conceptually frame the problem: each patient contributes a locally observed source-space patch that differs in both location and amplitude (scaling). Our goal is to stitch these patches together (quilting) while simultaneously adjusting their amplitudes to a common standard (mixed-effects modeling).

### 2.1 Materials

We analyzed stereoelectroencephalography (sEEG) and electrocorticography (ECoG) data from the **MNI Open iEEG Atlas**, as described by Frauscher et al. (2018). The dataset comprises recordings from 106 patients with therapy-refractory focal epilepsy, totaling 1,772 intracranial EEG (iEEG) electrodes.

Specifically, we utilized 60-second segments of resting-state wakefulness recordings (eyes closed). To balance temporal resolution with spectral estimation stability, the data were divided into 47 epochs of 2.5 seconds each, with a 50% overlap. All recordings were down-sampled to 200 Hz and bandpass filtered between 0.5 and 80 Hz, consistent with the preprocessing procedures in Frauscher et al. (2018).

For the sleep-stage validation, we additionally analyzed sleep recordings from the same dataset as described by von Ellenrieder et al. (2020). Sleep stages (N2, N3, and REM) were scored by board-certified epileptologists using concurrent scalp EEG according to the American Academy of Sleep Medicine (AASM) criteria (von Ellenrieder et al., 2020). Sixty-second artifact-free segments were selected from the first sleep cycle for each stage. The same channel inclusion criteria and preprocessing pipeline as for the wakefulness data were applied. Not all patients had usable recordings in every sleep stage, resulting in different sample sizes across conditions (Wake: *N* = 105; N2/N3: *N* = 87; REM: *N* = 61). To ensure that group comparisons were not confounded by differing electrode coverage, all statistical analyses between stages were restricted to patients with recordings available in both compared stages (i.e., paired analyses).

We adhered to the inclusion criteria established in the original study, retaining only channels that were: (1) located in non-lesional tissue as confirmed by MRI; (2) positioned outside the seizure onset zone; (3) free of interictal epileptic discharges and slow-wave anomalies; and (4) not recorded immediately following seizures or electrical stimulation. All electrode coordinates were mapped to the standard Montreal Neurological Institute (MNI) space to facilitate group-level aggregation. One subject was excluded from this study due to having only a single electrode, which was insufficient for the subsequent inverse problem-based scaling analysis.

### 2.2 Electrophysiological Source Imaging (ESI) Problem

For subject *i* (*i* = 1, …, *N*), the estimation of cortical activity from sensor data is framed as an inverse problem,

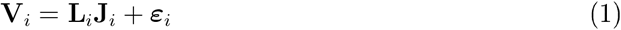

Here, 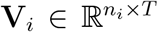 represents the sensor voltage with *n*_*i*_ sensor channels, and **J**_*i*_ ∈ ℝ^*M*×*T*^ represents the source current density with *M* source locations. 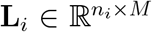 is the leadfield matrix quantifying the physical contribution of each source to each sensor. The estimated source activity 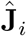 is obtained via eLORETA (Pascual-Marqui et al., 2011), which overcomes the depth bias inherent in algorithms like MNE and LORETA, exhibiting lower spatial dispersion and fewer ghost sources.

The leadfield matrix was computed using the DUNEuro Finite Element Method (FEM) via the Brainstorm toolbox (Medani et al., 2023) utilizing individual MRI-derived head models to account for tissue conductivity. Importantly, bipolar recordings were re-referenced to a unipolar montage to ensure compatibility with the leadfield calculation. This conversion is necessary because the leadfield calculation requires knowledge of the potential at each electrode with respect to a common reference, which is provided by the unipolar configuration.

### 2.3 Screening Method for High-Dimensional Regression

In the context of Electrical Source Imaging (ESI), estimating neural source activity from observed EEG signals constitutes an ill-posed and high-dimensional inverse problem. Given the sparse electrode coverage characteristic of sEEG and ECoG recordings, it is necessary to mitigate potential spurious source activity introduced by individual leadfields before the quilting procedure. To address this, we implemented a stable spatial projection solution for each leadfield by screening the individual source-space signals derived from ESI. Inspired by the marginal correlation screening of Fan and Lv (2008), we calculated the correlation coefficients between a large-scale set of simulated source signals and their reconstructed counterparts (obtained via the forward-inverse modeling loop using the individual lead eld). This metric was used to spatially select reliable source estimates, thereby establishing a stable projection relationship between the individual observations and the source space.

In the simulation, neural activity was constrained to the **cortical spatial manifold**. To produce source-space activity that conforms to the brain’s geometric topology, we generated spatially smooth null models based on the **geometric eigenmodes** (Fig. 2A) of the human cortex, using the following equation (Fig. 2B):

**Figure 2:**
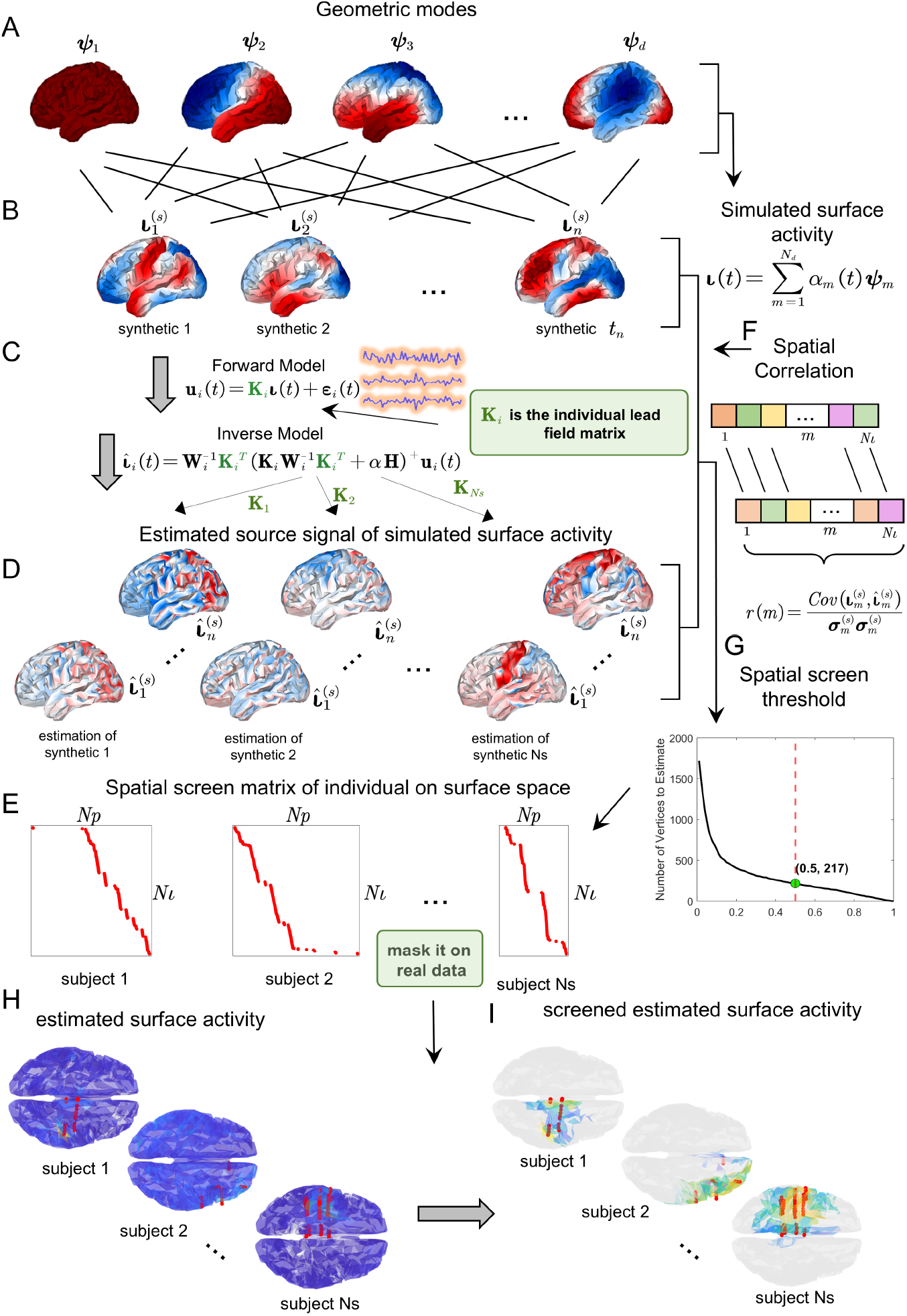
The procedure of spatial screening. A: Spatial geometric modes generated from the MNI-space cortical surface mesh (see Appendix A for details); B: Synthetic source activity 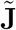 generated from snapshot *t*_1_ to *t*_*n*_ using Equation (2); C: Sensor samples 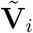 generated via the forward model based on individual leadfield matrices; D: Estimated source activity 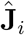 via the inverse model based on individual leadfield matrices; E: Spatial screening matrix corresponding to each leadfield; F: Schematic diagram of spatial correlation matrix computation; G: Threshold selection for spatial screening; H: Surface map of estimated activity 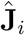 before screening; I: Surface map of estimated activity 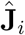 after applying screening masks.

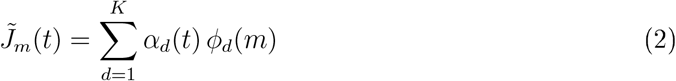

The simulated source activity at snapshot *t* is 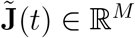, which represents a potential neural source configuration based on physiological assumptions. Here, *ϕ*_*d*_(*m*) is the *d*-th eigenfunction derived from the fs_LR_32k cortical surface template. We set *K* = 50 eigenmodes. A reconstruction analysis (Appendix B) confirmed that the screening correlation plateaus for *K* ≥ 40, with negligible improvement beyond *K* = 50, indicating that 50 eigenmodes suffice to probe all spatial directions resolvable by the forward-inverse loop. *α*_*d*_(*t*) is the *d*-th coefficient of snapshot *t*, generated via a Monte Carlo sampling algorithm. We generate synthetic source activity with *T* = 1000 sets of random coefficients 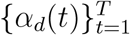.

To ensure a stable and accurate inverse projection for each leadfield, we assessed the spatial correlation between synthetic activity 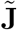 and estimated source activities 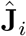. As shown in Fig. 2C, the estimated activity was obtained through a forward-inverse modeling loop utilizing the individual leadfield matrix.

The spatial correlation coefficient for subject *i* (with leadfield matrix **L**_*i*_), *r*_*i*_(*m*), between the simulated 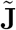 and reconstructed source activity 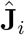 at location *m* is calculated as follows (illustrated in Fig. 2F):

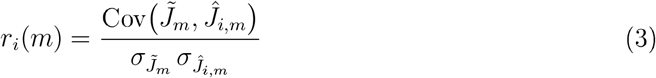

Here, 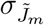 and 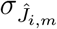 denote the standard deviations of 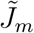 and 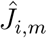, respectively. For subject *i*, the estimated source signal at location *m* is preserved when *r*_*i*_(*m*) > *τ*, where *τ* is the screening threshold (Fig. 2G), controlling the trade-off between spatial coverage and reconstruction fidelity. The threshold was selected via a systematic sensitivity analysis on synthetic data (Appendix C). This yields a reliable screening mask (Fig. 2E). By applying the obtained screening mask to the estimated source activity of the real data (Fig. 2H), we derive a reliable source estimate (Fig. 2I).

Here, it should be noted that the screening method based on correlation with a null model ensures spatial continuity inherited from the geometric eigenfunctions and guarantees the priority of higher leadfield gains, which cannot be achieved through global averaging methods like GSF.

### 2.4 Quilting with IOLMM

For each subject *i*, the screened source power is log-transformed to stabilize variance and convert multiplicative scaling differences into additive shifts:

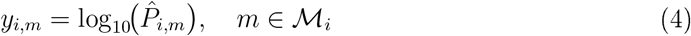

where 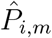 is the estimated source power spectral density at vertex *m*, and ℳ_*i*_ = {*m* : *r*_*i*_(*m*) > *τ*} denotes the set of vertices retained by the screening mask for subject *i*.

To construct a whole-brain source power map from sparse data, we aggregate the log-transformed, screened source power vectors 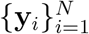 (Equation 4) across subjects using the IOLMM. The model explicitly accommodates two key sources of inter-subject variability: the spatial heterogeneity of electrode implantation and discrepancies in signal amplitudes.

For each subject *i* (*i* = 1, …, *N*), the log-transformed screened source power **y**_*i*_ (Equation 4) is modeled as:

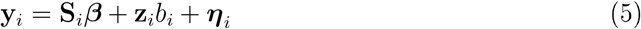

Because **y**_*i*_ is on the log_10_ scale, the inherently multiplicative amplitude scaling differences between subjects become additive components. The variables are:

- 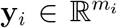 : Log-transformed screened source power for subject *i* (Equation 4), where *m*_*i*_ ≤ *M* is the number of vertices retained by the screening mask (*r*_*i*_(*m*) > *τ*).
- 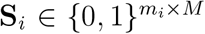 : Binary selection matrix constructed from the screening mask, mapping the full *M*-dimensional source space to the *m*_*i*_ retained vertices. Row *j* of **S**_*i*_ contains a single 1 at column *k* if *y*_*i,j*_ corresponds to cortical vertex *k*.
- ***β*** ∈ ℝ^*M*^ : Fixed effect vector representing the population-mean log source power across all *M* cortical vertices.
- 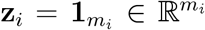 : Design vector for the random effect (a column of ones with length *m*_*i*_).
- 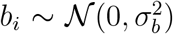: Subject-specific random effect (scalar), representing a global log-scale amplitude shift for subject *i*.
- 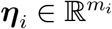 : Residual error vector, 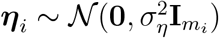.

Table 1 summarizes the physical interpretation of each model component.

**Table 1:** IOLMM effect structure and physical interpretation.

| Effect | Symbol | Interpretation | Estimated from |
| --- | --- | --- | --- |
| Fixed (group) | $\boldsymbol{\beta}$ | Population-mean log source power | Pooled screened sources |
| Random (subject) | $b_i$ | Subject-specific amplitude scaling | Individual log-scale offset |
| Residual | $\boldsymbol{\eta}_i$ | Measurement noise | Within-subject residual |

**Table 2:**
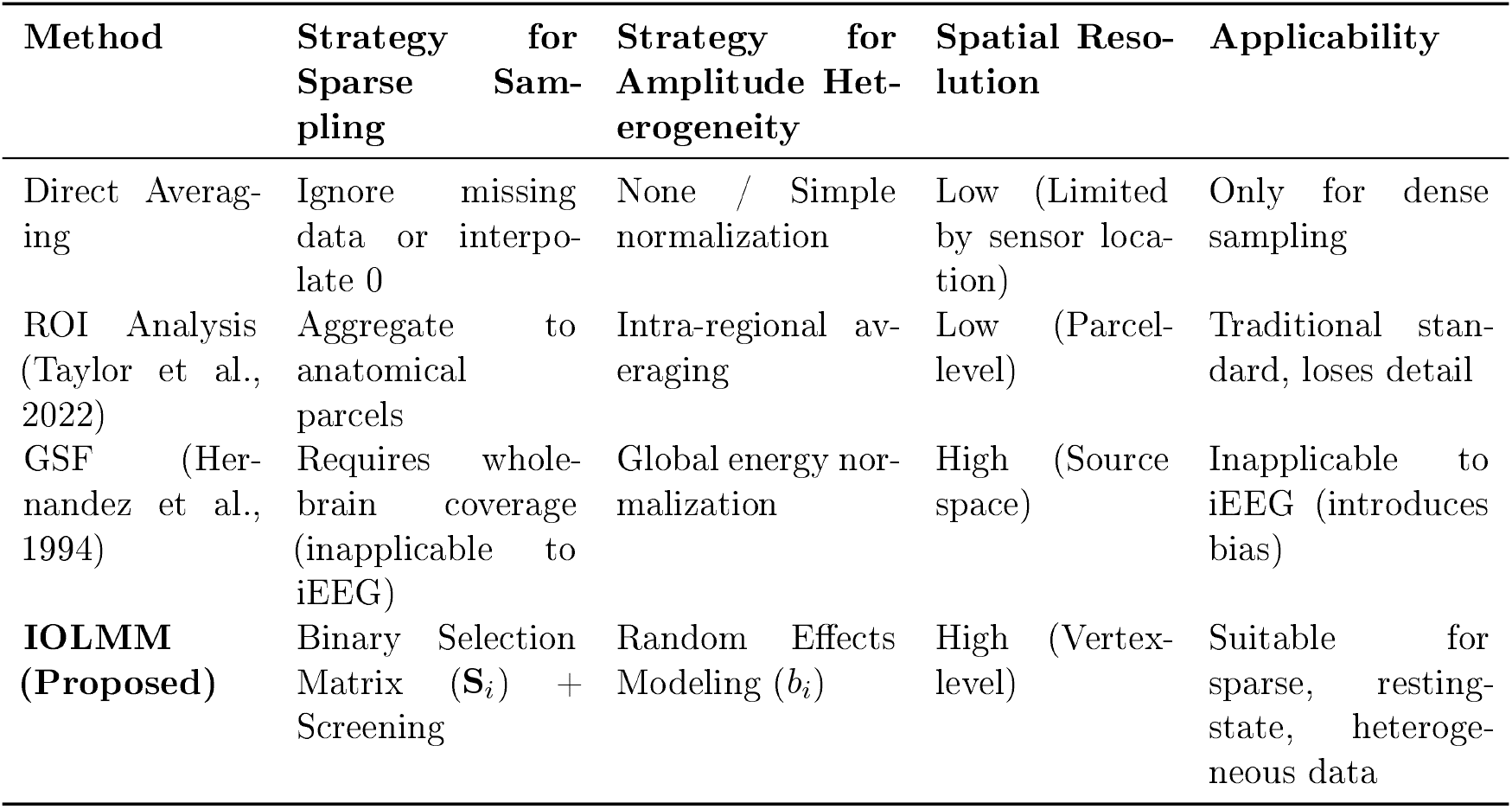
Comparison of methods for iEEG group analysis.

For a more general formulation, the residual covariance may assume structures beyond 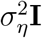, ranging from deterministic forms based on the Laplace Beltrami operator to Bayesian models employing a Wishart distribution (Gonzalez-Moreira et al., 2020; Paz-Linares et al., 2023). The parameters 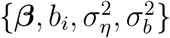 are estimated via the EM algorithm (Appendix D).

Once the EM algorithm (Appendix D) converges, the subject-specific scale factors are computed from the estimated random effects:

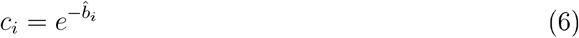

These scale factors adjust the amplitude of each subject’s source estimates to align them onto a common scale. Subtracting the random effect in log space yields the scale-adjusted observation:

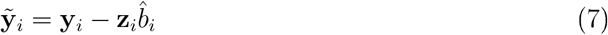

The group-level whole-brain activation map is then constructed by aggregating the adjusted source estimates across subjects. Under complete observation—i.e., **S**_*i*_ = **I**_*M*_ and *m*_*i*_ = *M* for all *i*–the estimator of ***β*** reduces to the simple sample mean:

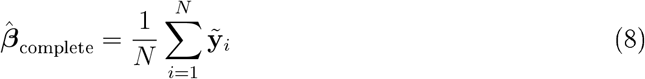

In practice, each subject observes only *m*_*i*_ ≪ *M* vertices, so the selection matrices **S**_*i*_ differ across subjects. A naïve application of Equation (8) to the back-projected vectors 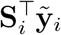 –i.e., dividing by *N*—would systematically underestimate vertices with low coverage: the *j*-th component would equal 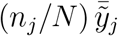, where *n*_*j*_ is the number of subjects observing vertex *j*. For vertices with *n*_*j*_ ≪ *N*, the factor *n*_*j*_*/N* attenuates the estimate toward zero. The EM-derived maximum-likelihood estimator (Appendix D) corrects for this by normalizing each vertex by its actual observation count:

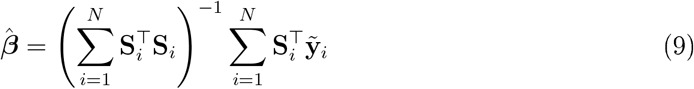

Since 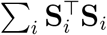 is diagonal with the *j*-th entry equal to *n*_*j*_, the *j*-th component of 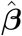 is the sample mean of 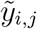 over only the *n*_*j*_ subjects who observe vertex *j*. When **S**_*i*_ = **I**_*M*_ for all *i*, every diagonal entry equals *N* and Equation (9) reduces to Equation (8).

Thus, the EM algorithm jointly estimates the per-subject scale factors *c*_*i*_ (Equation 6) and the group-level source power map 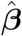 (Equation 9), tiling the **non-overlapping, partially covering**, and **scale-divergent** individual observations 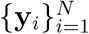 into a single whole-brain representation.

## 3 Results

To validate the effectiveness of the screening and quilting method for high-dimensional regression in selecting confident sources for ESI, we conducted a series of validation experiments based on simulated synthetic data. These simulations aimed to assess the method’s ability to accurately identify relevant sources and improve the reconstruction quality under conditions mimicking the sparse and non-uniform electrode coverage inherent in iEEG recordings. Here, we detail the protocol for generating synthetic samples, present the validation results for the screening and quilting steps, and summarize the biological validation of the framework using real iEEG data across sleep stages.

### 3.1 The Procedure of Synthetic Data Generation

To obtain realistic synthetic data, we employed structural geometric constraints for source data generation. Geometric modes (Fig. 3A) were generated based on the MNI-space cortical surface mesh (see Appendix A for details). The synthetic source activity was produced using Equation (2) (Fig. 3B), with coefficients derived via Monte Carlo sampling. We then calculated the source covariance matrix to facilitate the subsequent generation of sensor activity.

**Figure 3:**
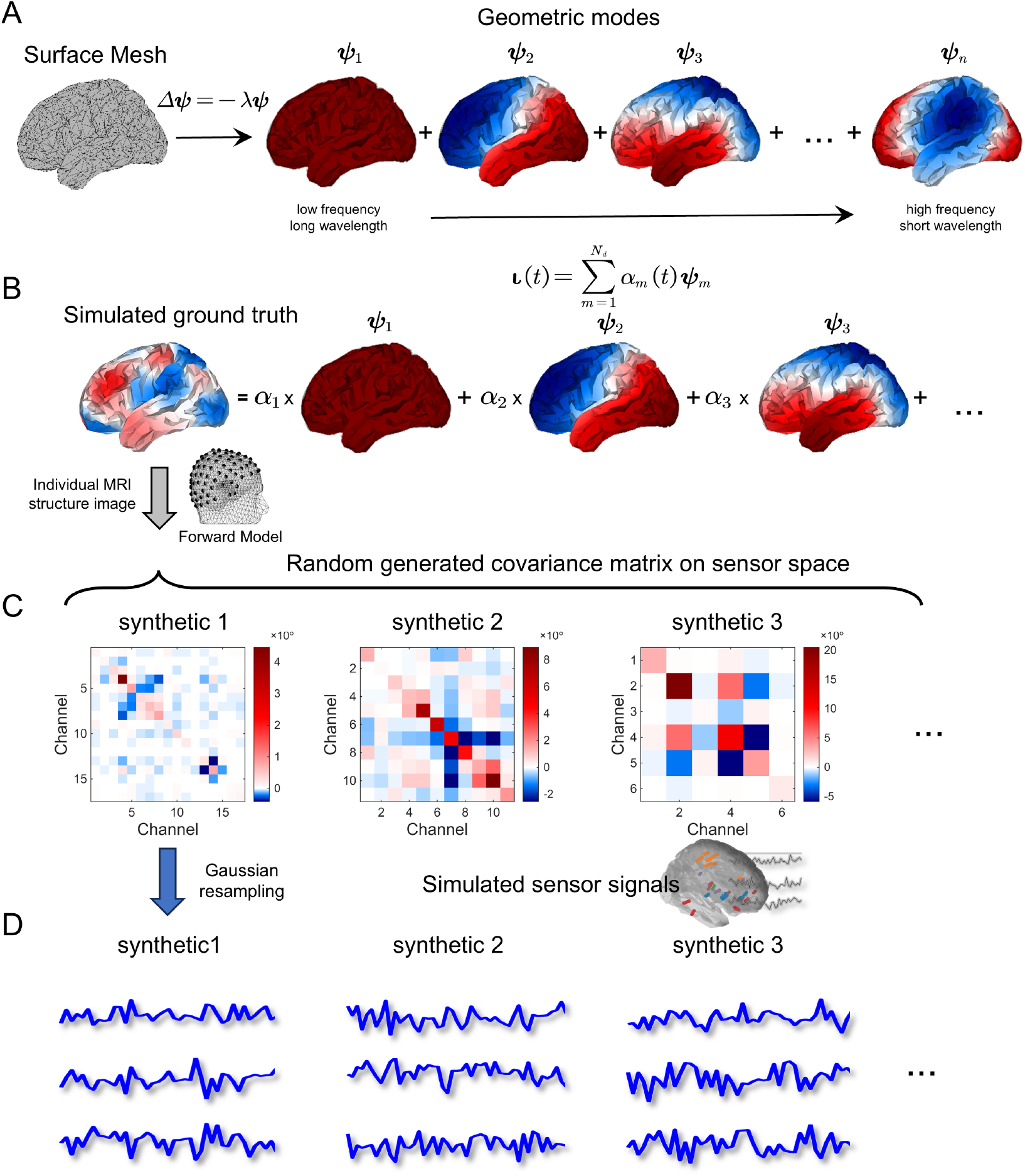
A: Spatial geometric modes obtained from the MNI-space cortical surface mesh (see Appendix A for details); B: Synthetic source activity generated with coefficients via Monte Carlo sampling; C: Covariance matrix in sensor space computed via the forward model for each individual leadfield matrix; D: Synthetic sensor signal sampled from a Gaussian distribution with the covariance matrix generated in C.

By applying the forward model with individual leadfield matrices, we obtained the covariance matrix in the sensor space (Fig. 3C). Consequently, subject-specific synthetic sensor activities were generated by sampling from a Gaussian distribution defined by this covariance 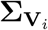 (Fig. 3D).

### 3.2 Validation of the Screening Method

To verify the necessity of spatial screening, we validated its e effectiveness using synthetic data by comparing the mean source activities obtained with and without screening against the ground truth. To eliminate potential confounding factors, the synthetic samples were generated with a unit amplitude scale.

Specifically, following the method described in Section 3.1, we generated synthetic sensor activities (Fig. 4A) and derived the Power Spectral Density (PSD) in the source space via inverse solving. For each sample, two strategies were applied: (1) estimation PSD without spatial screening 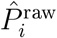 (Fig. 4B); and (2) estimation with spatial screening applied to the PSD 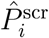 (Fig. 4C). We then calculated the geometric mean across samples for both resulting source PSDs (denoted as 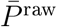 and 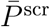). The correlation matrix (Fig. 4D) between the ground truth and the two mean maps reveals that the mean pattern derived with spatial screening maintains a high spatial correlation with the ground truth (*r* = 0.81). In contrast, the mean pattern without screening exhibits a correlation of only 0.54. These results strongly demonstrate the necessity of the spatial screening procedure.

**Figure 4:**
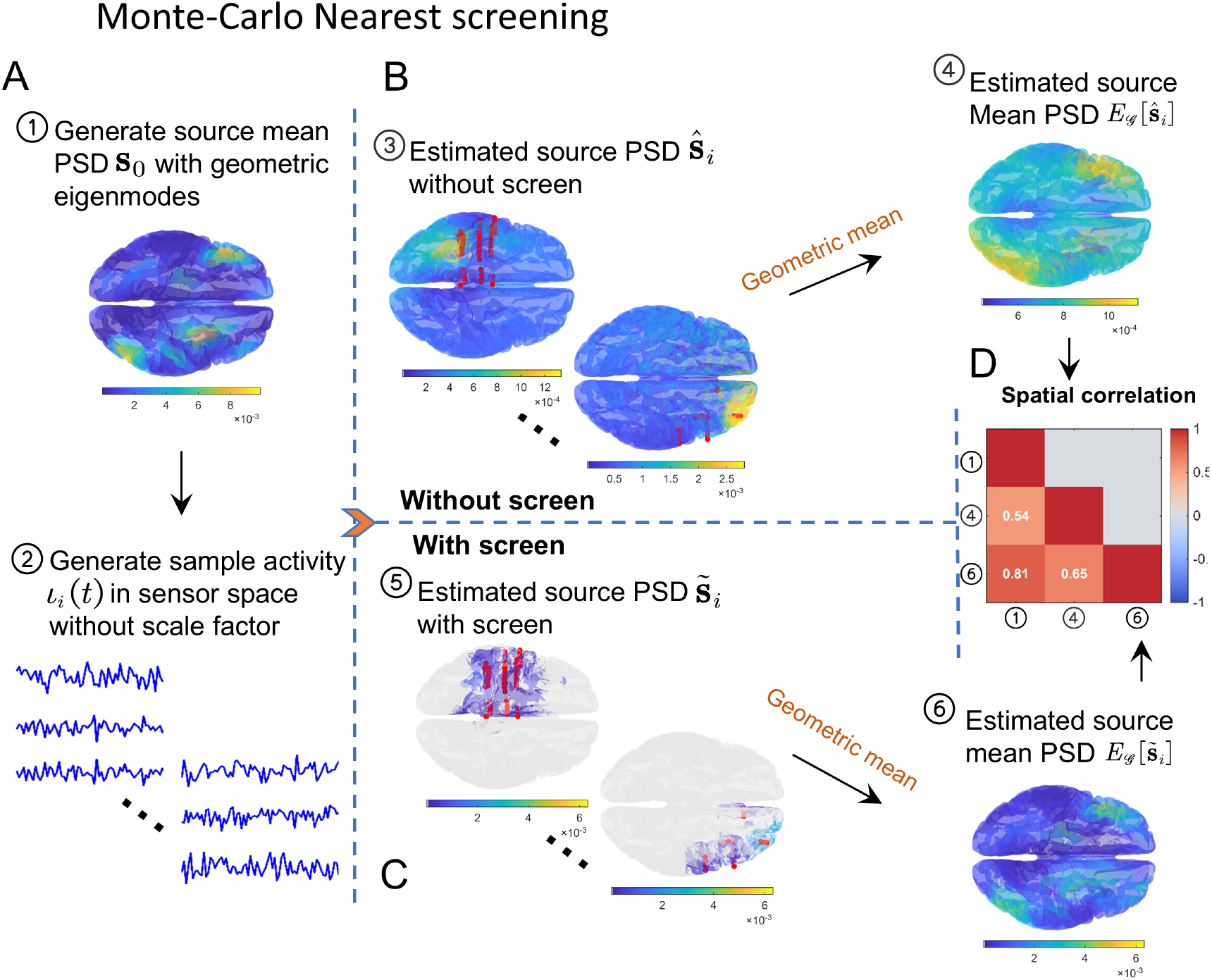
Validation of the spatial screening method. A: Ground truth source mean PSD 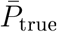 (left) and synthetic sensor activity 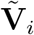 generated without scale factor (right); B: Estimated source PSD without screening 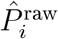 (left) and its geometric mean (right); C: Estimated source PSD with screening 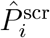 (left) and its geometric mean (right); D: Spatial correlation matrix between ground truth 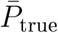, unscreened mean 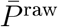, and screened mean 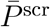.

### 3.3 Validation of the IOLMM Quilting Method

Here, we validated the effectiveness of scale correction across three different quilting methods using synthetic data. We compared the proposed IOLMM-based quilting method against two baselines: the straightforward Nearest Neighbor (NN) mean, and the Nadaraya-Watson (NW) interpolation algorithm. To obtain a systematic quantitative assessment, we employed three complementary metrics: (1) the Pearson correlation between the estimated source PSD and the ground truth PSD to quantify global pattern similarity; (2) ROC analysis for the detection of high-activity regions (top 25%, i.e., above the 75th percentile of ground truth values) to characterize local detection capability; and (3) the correlation between estimated and real scale factors, which provides direct evidence of scale estimation accuracy. As shown in Fig. 5A, the ground truth source mean map was generated using the method described in Section 3.1. The Nearest Neighbors quilting method represents the geometric mean across samples, while NW interpolation is a method that fills the source space directly from sensor space (details provided in Appendix E).

**Figure 5:**
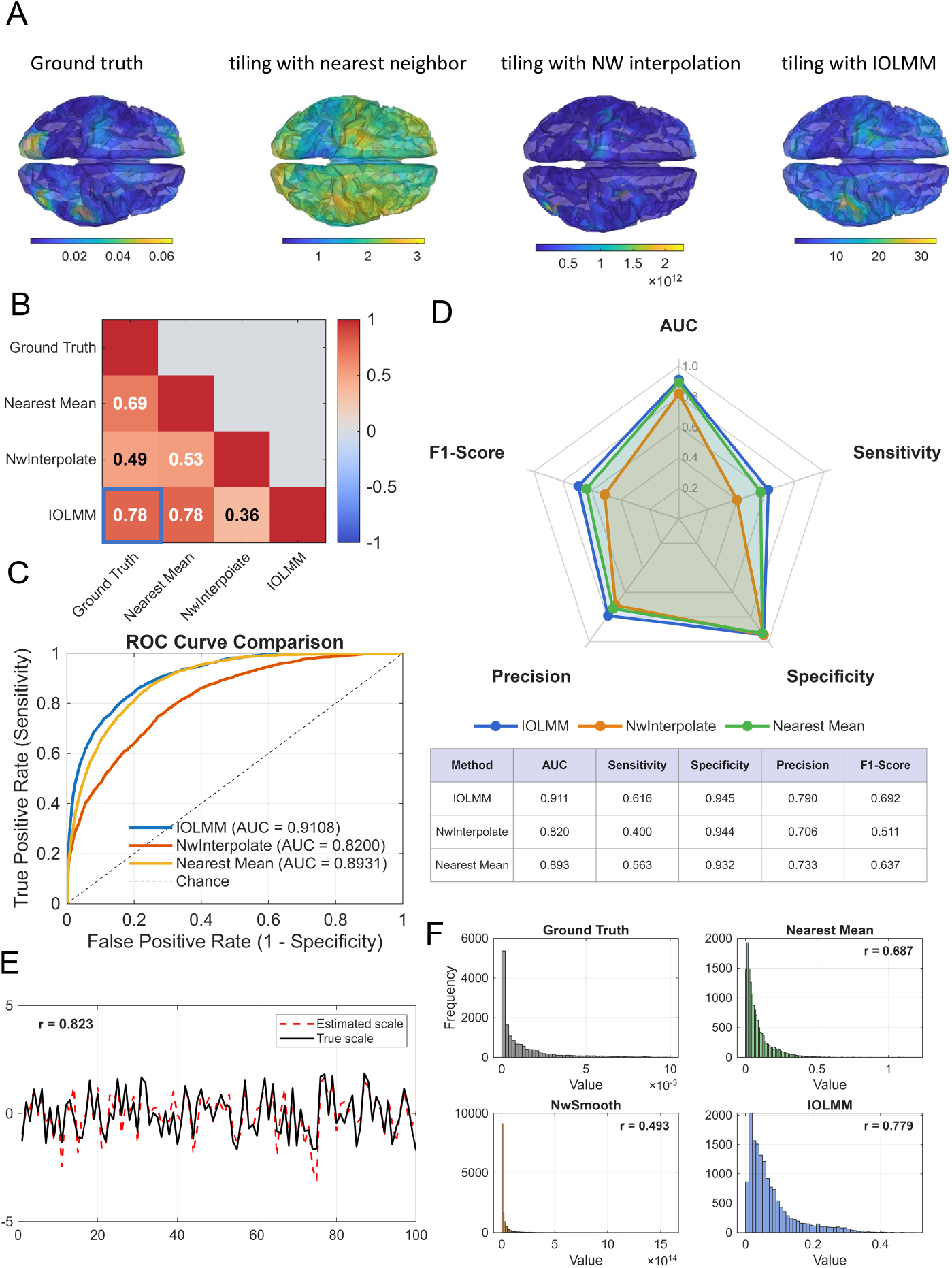
The validation of IOLMM. A: from left to right are: Generated ground truth source mean PSD, estimated source map which quilting with nearest neighbors, quilting with NW interpolation, quilting with IOLMM; B: Correlation matrix between ground truth with estimated source mean map; C: ROC curve of detection of cortical regions with high source power for IOLMM, NW interpolation and nearest mean methods; D: The classification performance of AUC, sensitivity, F1 score, precision, specificity among these three methods; E: The comparison between ground truth and estimated scale factor; F: The scale factors distribution comparison among methods.

The correlation matrix shows that the global spatial pattern derived from the IOLMM-corrected method is highly consistent with the ground truth, successfully recovering the overall activity distribution (*r* = 0.78, Fig. 5B).

Subject-specific amplitude scaling factors can severely impair the ability to localize original activity. Without correction, source signal intensity is overestimated in subjects with high scaling factors and underestimated in those with low scaling factors. This biases the group mean toward high-scale individuals, resulting in spatial pattern distortion. To further evaluate the spatial fidelity of the group map, we designed a detection task: identifying cortical regions with high source power (top 25th percentile of ground truth). The ROC analysis revealed that IOLMM-corrected estimates achieved a notably higher AUC (0.91) compared to Nearest Mean (0.89) and NW Interpolation (0.82) (Fig. 5C). This improvement demonstrates that removing subject-specific scaling factors enables the accurate recovery of the spatial distribution of source activity.

Furthermore, the classification radar chart (Fig. 5D) shows that the IOLMM correction outperformed other methods in terms of F1-score, AUC, sensitivity, specificity, and precision. These results directly verify that scale correction restores the capacity to localize active sources. The 75th percentile of ground truth values was used to define high-activity regions for the ROC analysis.

Finally, we compared the real sample scale factors with the estimated scale factors and their distribution. The results show that their correlation value is *r* = 0.823 (Fig. 5E), and the distribution correlation is 0.8 (Fig. 5F), which is higher than the other two methods, directly verifying the accuracy of IOLMM in scale estimation.

### 3.4 Biological Validation: Sleep Stage-Dependent Cortical Source Power

To demonstrate IOLMM on real data, we reconstructed whole-brain source power maps across four sleep stages (Wake, N2, N3, REM) from the MNI Open iEEG Atlas (Wake: *N* = 105; N2/N3: *N* = 87; REM: *N* = 61).

#### Stage-specific cortical power maps and lobe-level statistics

IOLMM-reconstructed source power maps (Fig. 6A) showed a progressive increase in cortical power from Wake through N2 to N3, with the strongest enhancement over frontal and cingulate cortex. REM power, by contrast, returned to or fell below Wake levels across most cortical regions. Rain-cloud plots of per-subject lobe-level source power (Fig. 6B) quantified this gradient: across all six lobes, the population ordering N3 > N2 > Wake > REM was maintained. FDR-corrected paired *t*-tests confirmed that N2 and N3 were significantly elevated relative to Wake in every lobe (all *p*_FDR_ *<* 0.001), while REM was significantly below Wake only in posterior lobes (parietal, occipital). The frontal lobe exhibited the largest dynamic range, spanning nearly one order of magnitude between N3 (−1.72) and Wake (−2.55) in log_10_ units, corroborating direct iEEG recordings of stage-dependent cortical power gradients (von Ellenrieder et al., 2020, 2022; Frauscher et al., 2018).

**Figure 6:**
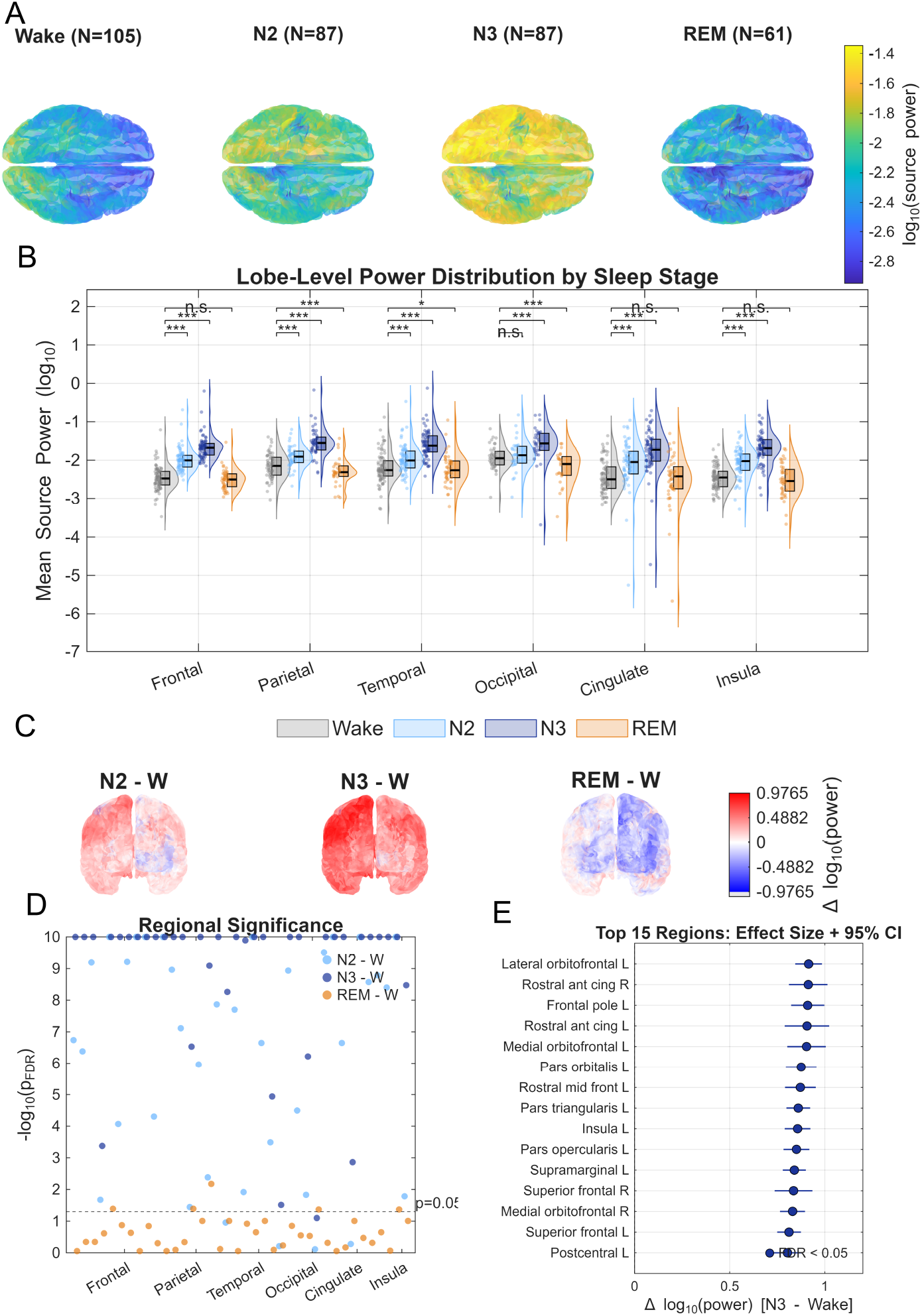
Sleep stage-dependent cortical source power reconstructed by IOLMM. A: Whole-brain source power maps for Wake, N2, N3, and REM; B: Raincloud plots of per-subject lobe-level source power; C: Vertex-wise difference maps (N2–Wake, N3–Wake, REM—Wake); D: Manhattan-style significance plot across DK-92 atlas regions; E: Forest plot of top 15 N3-vs-Wake regions ranked by effect size.

#### Spatial structure of stage contrasts and regional significance

Vertex-wise difference maps (Fig. 6C) revealed that N2–Wake and N3 –Wake differences were strongest over prefrontal and medial frontal cortex, while REM—Wake displayed a distinct anterior—posterior dissociation—consistent with the known antero-posterior power shift during REM sleep (Werth et al., 1996; De Gennaro et al., 2002). A Manhattan-style significance plot (Fig. 6D) across 72 DK-92 atlas regions showed that N3 vs. Wake reached significance in 67/72 regions (93%), N2 vs. Wake in 57/72 (79%), but REM vs. Wake in only 5/72 (7%)—mirroring the graded electrophysiological hierarchy. A forest plot of the top 15 N3-vs-Wake regions (Fig. 6E) showed that the largest effects (Δ log_10_ = 0.90−0.92) concentrated in prefrontal and anterior cingulate cortex (13/15 regions), consistent with the frontal predominance of slow-wave activity (Massimini et al., 2004; Bernardi et al., 2018).

## 4 Discussion

### 4.1 Quilting the Brain: IOLMM’s Solution to the Coverage Paradox

The central contribution of this study lies in formalizing and validating the **Incomplete Observation Linear Mixed-Effect Model (IOLMM)** as an effective solution to the “Quilting Problem” in intracranial EEG analysis. Rather than treating sparse coverage as a limitation to work around, IOLMM reframes it as a structured missingness problem amenable to principled statistical inference. Our results demonstrate that this formulation can reconstruct coherent, whole-brain source power maps that recover known physiological patterns—including the frontal predominance of slow-wave activity and the graded NREM hierarchy—from the fragmented observations of individual patients. This outcome suggests that the coverage paradox, while a genuine obstacle for channel-level analyses, is not a fundamental barrier when the problem is cast in source space with appropriate hierarchical modeling.

Key to the success of IOLMM is its handling of **Scaling Ambiguity**. As discussed in the introduction, the traditional Global Scale Factor (GSF) fails in iEEG because it conflates the extent of observation with signal magnitude. Our analysis further confirms that when sensor arrays sample only a fraction of the brain volume, the assumption of total energy conservation underlying GSF is broken. IOLMM effectively decouples the biological signal of interest (i.e., the fixed effect, ***β***) from confounding variations introduced by electrode impedance, amplifier gain differences, and, crucially, coverage density differences, by modeling subject-specific scaling factors (*b*_*i*_) as **Random Effects** within a mixed-model framework.

The logarithmic transformation employed in the model converts multiplicative scaling differences into additive biases, which are then efficiently estimated via the EM algorithm. This mathematical treatment is not merely for computational convenience but deeply reflects the physical nature of the signal—gain factors physically act multiplicatively on neural current density. The resulting group-level source power map 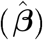 represents a “consensus” physiology—a statistical estimate of the expected iEEG power distribution if one could record from all positions simultaneously in a standard subject. This establishes a direct bridge between the sparse world of highdelity iEEG and the dense, continuous atlases of fMRI/MEG.

It is instructive to compare IOLMM with existing approaches to iEEG normative mapping. Frauscher et al. (2018) constructed the first normative iEEG atlas using channel-level spectral features aggregated into anatomical parcels, while Taylor et al. (2022) extended this to normative z-score mapping for epileptogenic tissue localization. Both approaches operate in *sensor space*, inheriting the spatial resolution of the electrode array itself. In contrast, IOLMM operates in *source space* on the cortical surface mesh, offering vertex-level resolution that is independent of individual electrode placement. Furthermore, neither prior approach explicitly models inter-subject amplitude scaling; they rely on ROI averaging or simple z-scoring that can be biased by heterogeneous electrode coverage. IOLMM addresses this through random-effects modeling of subject-specific scaling factors, making it better suited for structurally incomplete data.

### 4.2 The Necessity of High-Dimensional Screening and the Value of Geometric Constraints

Another key finding of this study is that rigorous source space screening is indispensable prior to “quilting”. The iEEG inverse problem is notoriously unstable for regions distant from electrode contacts. Without screening, ESI algorithms (such as eLORETA) generate substantial “Ghost Sources” in uncovered regions, often driven by noise or volume conduction effects from active areas. Simply averaging these noisy estimates into a group atlas would severely degrade spatial resolution and introduce false positives, negating the high spatial precision advantage of iEEG.

We addressed this by drawing on the concept of “Sure Independence Screening” (SIS) from high-dimensional regression (Fan and Lv, 2008). By simulating smooth source activity based on the brain’s **Geometric Eigenmodes**, we created a biologically plausible “null model”. Calculating the correlation between simulated sources and their inverse problem reconstructions provides a reliable metric. This process effectively generates a “Confidence Mask” for each subject, precisely identifying the cortical patches where that subject’s electrode array provides credible information.

This screening process transforms the input for IOLMM from a dense but noise-filled matrix into a sparse but high-confidence matrix. IOLMM then acts as the “weaver,” stitching these high-confidence patches together. This “screen first, quilt second” strategy ensures that the final group atlas retains the high spatial frequency information inherent to iEEG signals, rather than smoothing it away through low-confidence extrapolation. This echoes recent theoretical work on “Graph Quilting,” which aims to infer the global structure of graph models from block-missing covariance data (Vinci et al., 2023). Although our approach focuses on activation mapping (first-order statistics) rather than connectivity (second-order statistics), the underlying logic is consistent: global inference requires identifying and exploiting overlapping information within reliable data subspaces.

An important property of the screening procedure is that the per-vertex reliability metric *r*_*i*_(*m*) (Equation 3) is determined by the **resolution matrix R**_*i*_ = **T**_*i*_**L**_*i*_, which encodes how faithfully each cortical vertex survives the forward—inverse modeling loop for a given electrode configuration. The geometric eigenmodes serve only as spatially smooth probe signals; any sufficiently smooth and spatially distributed basis (e.g., spherical harmonics, Fourier basis) would yield similar screening masks, because the discriminating factor is the subject-specific forward—inverse loop, not the input signal’s spectral composition. This con rms that screening is a data-independent preprocessing step whose selectivity is governed by electrode geometry, not by the choice of probe basis.

The following table summarizes the comparison between the IOLMM method and traditional approaches in handling iEEG group analysis challenges:

### 4.3 Construction of Normative Atlases and Clinical Translational Significance

Generating reliable group-level source power maps has direct implications for developing **Normative Atlases** for iEEG. Prior work, such as the seminal study by Frauscher et al. (2018), relied primarily on strict exclusion criteria and simpler aggregation methods to define “normal” iEEG activity. While foundational, these atlases mostly utilized channel-based or ROI-based metrics. Our source-space method offers a high-resolution alternative defined continuously on the cortical manifold.

By providing estimates of vertex-level mean spectral power and its total variance (captured by residual error 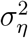 and random effects variance 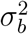), IOLMM enables the calculation of **Z-score maps** for individual patients relative to a normative baseline. Clinically, this function has practical value. In presurgical epilepsy evaluation, identifying the seizure onset zone often relies on detecting subtle deviations in background activity or interictal discharges. A normative atlas allows clinicians to statistically quantify the “degree of abnormality” at the source level, potentially highlighting Focal Cortical Dysplasia (FCD) or other lesions that are subtle on MRI but spectrally distinct.

Furthermore, this method facilitates the fusion of iEEG with other modalities. The “quilted” iEEG atlas can be directly correlated with fMRI BOLD signals or MEG source reconstructions in a common space (e.g., MNI or fs_LR_32k). This promotes cross-modal validation studies, such as using the temporal precision of iEEG to dissociate neural drivers of hemodynamic signals observed in fMRI, or validating the source localization accuracy of non-invasive MEG.

### 4.4 Biological Validation: Sleep Stage-Dependent Source Power

The sleep-stage application validates the IOLMM pipeline by demonstrating that known physiological patterns can be recovered from fragmented iEEG coverage. Despite each subject contributing electrodes covering only a small fraction of the cortex, IOLMM recovered the frontal predominance of NREM slow-wave activity—a topographic signature established by high-density scalp EEG (Werth et al., 1996; Massimini et al., 2004) and confirmed by direct cortical recordings (von Ellenrieder et al., 2020). That the same spatial pattern emerges from quilting ~100 partial-coverage subjects provides evidence that the screening step effectively removes ghost sources and the IOLMM quilting correctly adjusts inter-subject amplitude scaling.

The method also exhibits appropriate sensitivity to graded electrophysiological effects: N3 effects were detected in 93% of regions, N2 in 79%, REM in only 7%, mirroring the known hierarchy and suggesting that IOLMM output retains sufficient signal-to-noise for standard parametric inference. The near-absence of significant REM effects rules out systematic inflation bias. Furthermore, the raincloud plots show that per-subject distributions overlap across stages within each lobe while population medians maintain strict ordering—a property required for normative Z-score atlases where individual patient deviations must be meaningfully quantified.

### 4.5 Limitations and Methodological Considerations

While the IOLMM framework shows considerable potential, several limitations warrant further discussion.

First, the effectiveness of “quilting” relies heavily on a core assumption: that the **Fixed Effect** (i.e., group mean) is a meaningful biological entity. This implicitly assumes cross-subject **Functional Homology**–that after stereotactic normalization, the “Fusiform Face Area” or “Default Mode Network” resides in roughly the same anatomical location across individuals. Although cortex-based alignment improves this correspondence compared to Volume-based registration, individual differences in functional topology may contribute residual variance. IOLMM accounts for **amplitude** variation via random effects but does not explicitly model spatial variation (i.e., displacement of functional peaks). Future work could incorporate **spatial** random effects or Optimal Transport metrics to further reduce this residual.

A consideration shared by all iEEG normative studies is that the data originate from patients with epilepsy rather than healthy volunteers. This practice is well-established and defensible (Rutherford et al., 2023): rigorous channel exclusion criteria—removing channels within the seizure onset zone, those exhibiting interictal epileptic discharges, and those in lesional tissue (Frauscher et al., 2018) –yield recordings that approximate normal cortical activity. Moreover, epilepsy is increasingly recognized as a network disorder, and recent evidence shows that more complete resection of seizure onset regions is not necessarily associated with more favorable surgical outcomes (Gascoigne et al., 2024), suggesting that the boundary between “pathological” and “normal” tissue is itself uncertain. Nevertheless, electrode placement is determined by clinical need rather than scientific design, so frequently implanted regions (e.g., the temporal lobe) are over-represented relative to less commonly targeted areas. IOLMM handles the structural pattern of missingness via the selection matrix **S**_*i*_, but the statistical power of the group atlas is necessarily spatially non-uniform, with higher confidence in densely sampled regions. Future multi-center data collection efforts could aim to broaden spatial coverage by pooling cohorts with complementary implantation strategies.

Additionally, the current sleep-stage analysis is restricted to broadband source power. This broadband characterization provides the foundational validation of the quilting pipeline; frequency-resolved maps (e.g., slow oscillation vs. sigma/spindle band) are a natural next step that would yield more specific physiological fingerprints.

Furthermore, eLORETA source localization, while exhibiting zero localization error for point sources, introduces spatial blurring (effective resolution ~15−20 mm) that propagates into the IOLMM vertex-level estimates. The geometric eigenmode screening mitigates ghost sources from the inverse solver, but cannot recover spatial detail finer than the leadfield resolution. Consequently, IOLMM maps should be interpreted at the gyral scale rather than at individual cortical columns.

Looking ahead, the framework opens several concrete avenues. A direct extension is applying IOLMM to **time-frequency data**, enabling the quilting of event-related spectral dynamics or dynamic functional connectivity. This would require expanding the scalar random effect *b*_*i*_ into time-varying or frequency-dependent scaling functions. Comparing IOLMM with other data **harmonization** techniques (e.g., ComBat; Johnson et al., 2007) is also warranted. While ComBat excels at removing batch effects in dense neuroimaging data, its application to sparse, structurally missing data is not straightforward; hybrid methods combining ComBat’s empirical Bayes estimation with IOLMM’s mixed-model structure may yield further improvements in multi-center harmonization. Finally, the quilted atlas could serve as a reference for training generative models that fuse sparse iEEG with dense fMRI or MEG data, leveraging the complementary spatial and temporal resolution of these modalities.

### 4.6 Conclusion

In summary, the “quilting” of multi-center intracranial EEG data represents a critical step toward constructing a high-resolution human brain functional atlas. By combining high-confidence source screening based on geometric eigenmodes with Incomplete Observation Linear Mixed-Effect Models, we provide a mathematically principled method to overcome the obstacles of sparse sampling and heterogeneous scaling. This approach not only validates the feasibility of group-level inference using iEEG but also establishes a new standard for building normative electrophysiological atlases, thereby bridging the chasm between neurons and networks, and between individual patients and the population.

## Ethics Statement

This study used retrospective, de-identified data from the MNI Open iEEG Atlas (Frauscher et al., 2018). All patients provided written informed consent for their clinical data to be used for research purposes, and the original study was approved by the Research Ethics Board of the Montreal Neurological Institute and Hospital (McGill University). No additional ethical approval was required for the present secondary analysis of the publicly available, de-identified dataset.

## Data Availability

The MNI Open iEEG Atlas data used in this study are publicly available at https://mni-open-ieegatlas.research.mcgill.ca. The IOLMM code is available at https://github.com/ccclab-bigdata/IOLMM.

## CRediT Author Statement

M.L., Y.W., and P.A.V-S. jointly conceived and designed the study. Y.W. and M.L. curated the data, performed the whole data analyses, and wrote the manuscript. M.L.B-V. contributed to the development and critical revision of the manuscript.

## Acknowledgements

We thank Dr. Nicolás von Ellenrieder for generously sharing the MNI iEEG Atlas dataset and for his expert guidance on its use. This work was supported by the National Key R&D Program of China (2024YFE0215100), the CNS Program of the University of Electronic Science and Technology of China (UESTC) (Grant No. Y03023206100204). M.L. was supported by Hangzhou Dianzi University Seed Fund Project (KYS055623037) and Zhejiang Provincial Higher Education Institutions’ Basic Operations Project (GK239909299001-025).

## A Calculation of Geometric Eigenfunctions

To obtain the eigenmodes, we defined the cortex space using a triangular cortical surface mesh obtained by downsampling the FreeSurfer cortical reconstruction of the ICBM152 template via Brainstorm (Tadel et al., 2011). Two resolutions are used: 2,003 vertices for screening validation and 15,002 vertices for the final empirical atlas. As this template is a public standard and independent of our data sample, it avoids any data information leakage. We then constructed the Laplace Beltrami operator (LBO) from this surface mesh, which captures local vertex-to-vertex spatial relations and curvature. By solving the eigenvalue problem of the following equation, we can obtain the geometric eigenmodes,

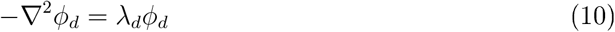

where ∇^2^ is the Laplace-Beltrami operator. 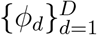 are the eigenmodes with eigenvalues 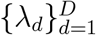. The eigenvalues are sorted in ascending order and all eigenfunctions are mutually orthogonal, forming a complete geometric basis. Synthetic source activity was generated as a linear combination of the first *K* = 50 eigenmodes (see Appendix B for the sensitivity analysis justifying this choice).

## B Eigenmode Count Sensitivity Analysis

To verify the choice of *K* = 50 eigenmodes for the screening probe signals (Equation 2), we assessed how the number of eigenmodes affects the fidelity of the forward–inverse reconstruction loop. For a representative sEEG electrode configuration, we generated a smooth ground-truth source pattern **s** = **Φ**_1:200_ **c** using all 200 eigenmodes with random coefficients **c**. We then truncated the basis to the first *K* modes (*K* ∈ {5, 10, 15, 20, 30, 40, 50, 75, 100, 150, 200}), passed the truncated signal through the resolution matrix **R**_*i*_ = **T**_*i*_**L**_*i*_, and measured the Pearson correlation between the truncated reconstruction **R**_*i*_**Φ**_1:*K*_ **c**_1:*K*_ and the full reconstruction **R**_*i*_ **s**.

The reconstruction correlation increased from *r* = 0.63 at *K* = 5 to *r* = 0.84 at *K* = 40, after which it plateaued: *r* = 0.84 at *K* = 50 and *r* = 0.86 at *K* = 100 (Fig. 7). This plateau indicates that beyond *K* ≈ 40, additional eigenmodes contribute spatial frequencies that the resolution matrix cannot resolve, yielding diminishing returns for screening. We therefore adopted *K* = 50 as a conservative choice within the plateau region.

**Figure 7:**
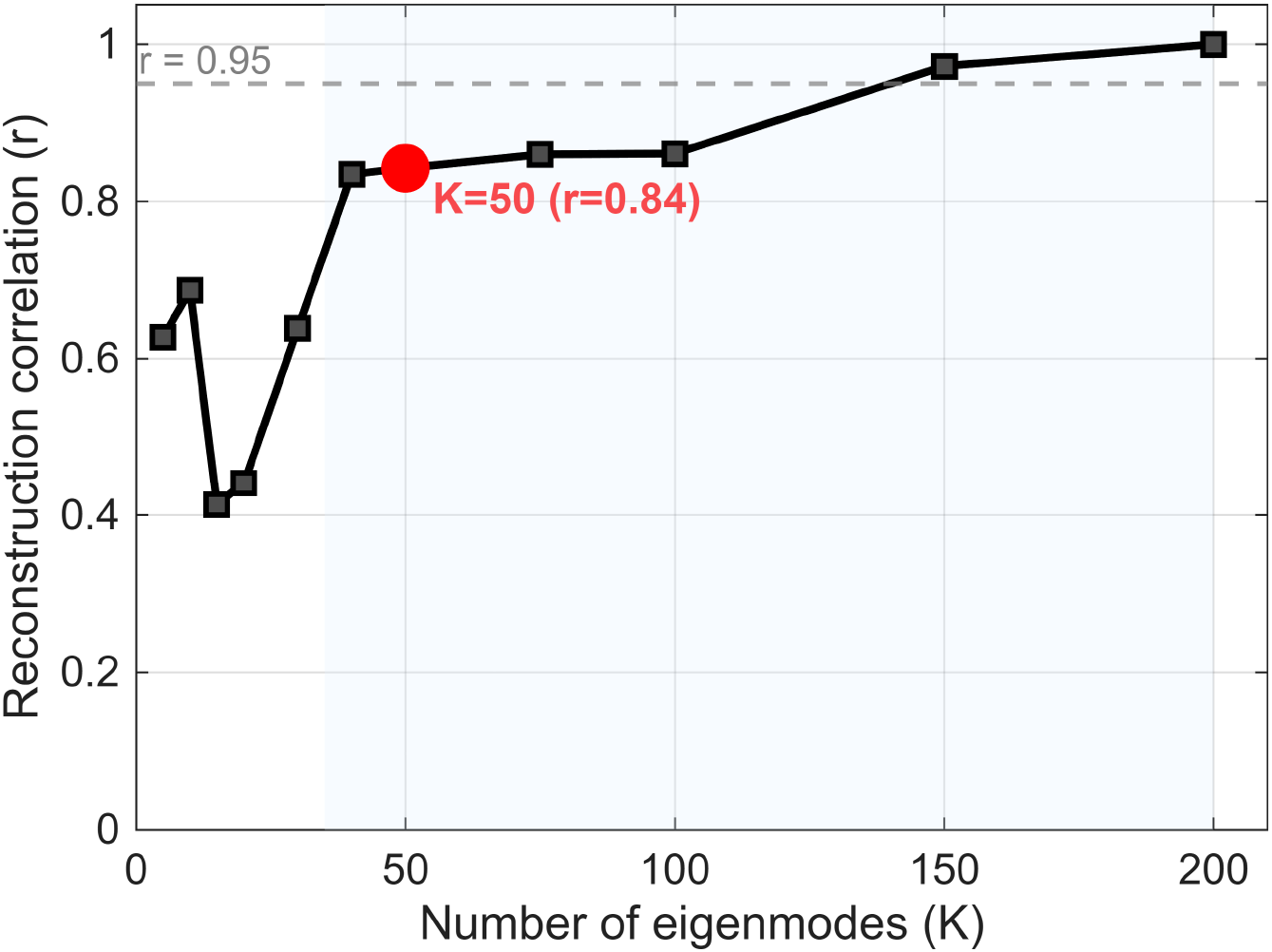
Reconstruction correlation as a function of eigenmode count *K*. A smooth ground-truth source pattern (generated from 200 eigenmodes) was truncated to the first *K* modes and passed through the resolution matrix **R**_*i*_ = **T**_*i*_**L**_*i*_ of a representative sEEG electrode configuration. The red dot marks *K* = 50 (*r* = 0.84), which lies within the plateau region (*K* ≥ 40). The shaded area indicates the plateau region where reconstruction quality is stable.

## C Screening Threshold Sensitivity Analysis

The screening procedure retains source vertices where the spatial correlation *r*_*i*_(*m*) exceeds a threshold *τ*, controlling the trade-off between spatial coverage and reconstruction delity. A lenient threshold retains many vertices, including unreliable “ghost sources”, whereas a stringent threshold keeps only high-fidelity vertices but reduces coverage, potentially leaving gaps in the quilted map. To select *τ*, we performed a systematic sensitivity analysis on synthetic data (see Section 3.1 for details). We swept *τ* across [0.1, 0.9] with a step size of 0.05 (17 values). For each *τ*, we executed the complete pipeline—screening, IOLMM fitting (EM algorithm), and quilting—and evaluated reconstruction quality by computing the Pearson correlation between the quilted source map 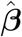 and the synthetic ground truth **s**_0_, as well as the area under the ROC curve (AUC) for detecting high-activity regions (top 25th percentile). Two baseline methods (Nearest Neighbor mean and Nadaraya-Watson interpolation) were evaluated in parallel to assess whether method ranking was consistent across threshold values. We defined the optimal threshold *τ*^∗^ as the smallest *τ* at which the IOLMM reconstruction correlation reached 95% of its maximum value across all tested thresholds, balancing reconstruction quality against coverage.

Mean vertex coverage decreased monotonically from 32.7% (*τ* = 0.1) to 2.0% (*τ* = 0.9), confirming that stricter thresholds progressively exclude more source locations (blue curves in Fig. 8A and B). The IOLMM reconstruction quality increased monotonically with *τ*, as measured by both Pearson correlation (Fig. 8A; *r*: 0.63 at *τ* = 0.1 to 0.88 at *τ* = 0.85) and AUC (Fig. 8B; 0.83 to 0.95). This demonstrates that the linear mixed-effect quilting mechanism effectively recovers the full source map even from sparse, high-quality inputs. In contrast, the Nearest Neighbor mean exhibited a non-monotonic pattern, peaking at *τ* ≈ 0.50 (*r* = 0.69) before declining at stricter thresholds due to insufficient spatial sampling. The Nadaraya—Watson interpolation, which operates directly in sensor space, was largely invariant to *τ* (*r* ≈ 0.50), serving as a *τ*-independent baseline.

**Figure 8:**
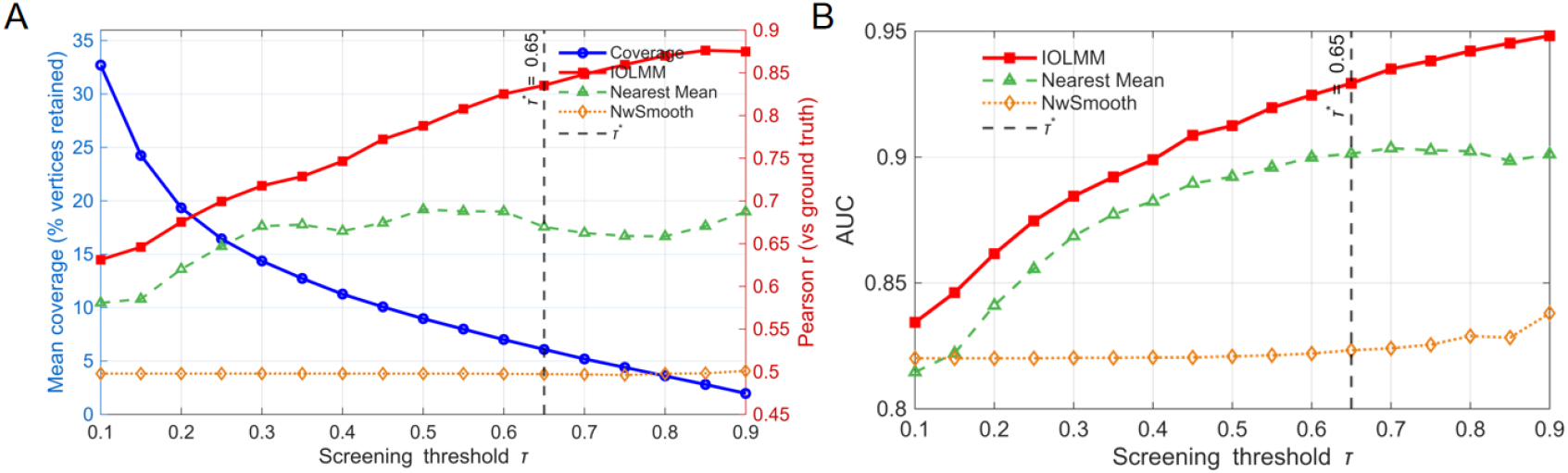
Sensitivity of reconstruction quality to screening threshold *τ*. (A) Coverage –correlation trade-off. Left y-axis (blue circles): mean fraction of cortical vertices retained per subject. Right y-axis: Pearson correlation between the quilted source map and synthetic ground truth for IOLMM (red squares), Nearest Neighbor mean (green triangles), and Nadaraya–Watson interpolation (orange diamonds). (B) Coverage—AUC trade-off. Same layout as (A), with the right y-axis showing area under the ROC curve for detecting high-activity cortical regions (top 25th percentile). In both panels, the dashed vertical line indicates the selected threshold *τ*^∗^ = 0.65, defined as the smallest *τ* at which the IOLMM metric reaches 95% of its maximum value. The method ranking (IOLMM > NN > NW) is consistent across all tested thresholds, confirming robustness to this parameter choice.

Applying the 95%-of-maximum criterion yielded *τ*^∗^ = 0.65, at which the IOLMM achieved *r* = 0.84 and AUC = 0.93 with a mean coverage of 6.1% of cortical vertices per subject. Crucially, the method ranking (IOLMM > Nearest Mean > NW interpolation) was consistent across all 17 tested *τ* values (100%), indicating that the comparative conclusions reported in this study are robust to threshold selection (Fig. 8).

## D EM Algorithm for the IOLMM

### EM Algorithm Steps

1. **Initialization:** Initialize the fixed effect ***β***^(0)^ and set the stop criteria. Initialize the variance components 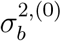 with a random value smaller than 1, 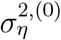.
2. **Expectation Step (E-step)** (for each subject): Compute the conditional moments of the missing random effect *b*_*i*_ given the current estimates of the fixed effect and variance components 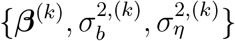 for each subject *i* with the following formulas,

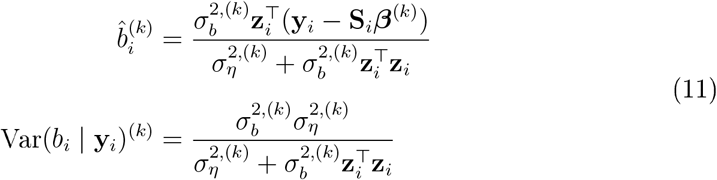
3. **Maximization Step (M-step)** (for global likelihood): Update the estimates of the fixed effect and variance components by maximizing the expected log-likelihood computed in the E-step. **Fixed Effect Update:**

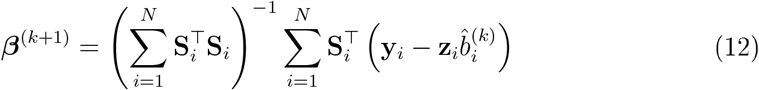

This is the estimator presented in Equation (9), applied iteratively within the EM loop. **Variance Components Update:** Update 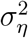 and 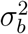 based on the residuals and estimated random effects.

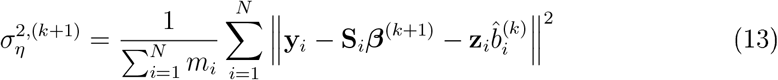

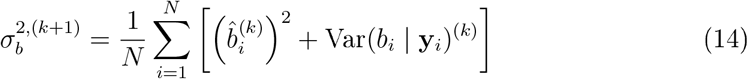
4. **Convergence Check:** Evaluate the relative change in the log-likelihood between iterations. The algorithm terminates when the relative change falls below 10^−6^ or the number of iterations reaches 1,000, whichever occurs first.
5. **Iteration:** Repeat the E-step and M-step until convergence.

## E Nadaraya-Watson Kernel Regression for Source Space Interpolation

As a baseline quilting method, we employ the Nadaraya-Watson (NW) kernel regression estimator to interpolate source power from sparse sensor-level observations onto the dense cortical surface mesh. Unlike the IOLMM, which explicitly models subject-specific scaling via random effects, NW interpolation is a non-parametric approach that constructs a continuous surface map by spatially weighted averaging of sensor-level power estimates.

For each subject *i*, let 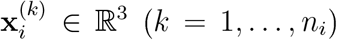 denote the 3D coordinates of the *n*_*i*_ electrode midpoints (in MNI space), and let 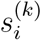 denote the corresponding sensor-level power estimate (e.g., 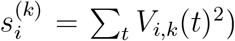). The goal is to estimate the source power 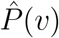 at every vertex *v* ∈ ℝ^3^ of the cortical surface mesh.

We adopt an isotropic Gaussian kernel. To ensure scale-invariance across coordinate dimensions, all electrode and vertex coordinates are first standardized (z-scored) using the electrode population statistics:

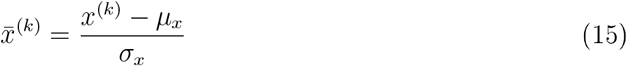

where *µ*_*x*_ and *σ*_*x*_ are the mean and standard deviation computed across all electrode positions from all subjects.

The Gaussian kernel weight between electrode position 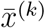 and query vertex 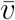 is defined as:

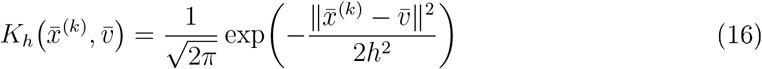

where *h* > 0 is the bandwidth parameter controlling the spatial extent of the smoothing.

The NW estimator computes the source power at each cortical vertex *v* as a kernel-weighted average of all sensor-level power observations:

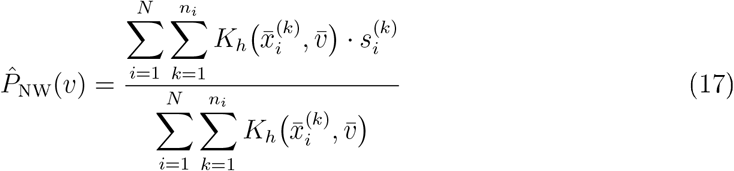

This formula pools all electrode observations from all subjects, weighting each observation by its spatial proximity to the query vertex. Electrodes closer to vertex *v* contribute more to the estimate than distant ones.

The bandwidth *h* controls the bias-variance trade-off : a small *h* yields a locally adaptive but noisy estimate, while a large *h* produces a smoother but potentially over-smoothed map. We select *h* using Generalized Cross-Validation (GCV), which provides a data-driven, leave-one-out-equivalent criterion without requiring explicit cross-validation loops.

The NW estimator can be expressed in matrix form. Let **s** = (*s*_1_, …, *s*_*n*_)^⊤^ denote the pooled sensor power vector across all 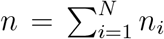 electrodes from all subjects. The NW estimate at the electrode locations themselves is:

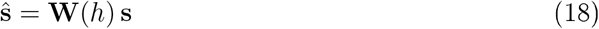

where the smoother matrix **W**(*h*) ∈ ℝ^*n*×*n*^ has entries:

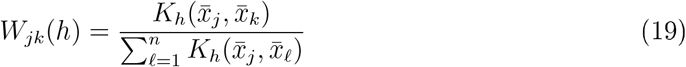

The GCV criterion selects the bandwidth *h*^∗^ that minimizes:

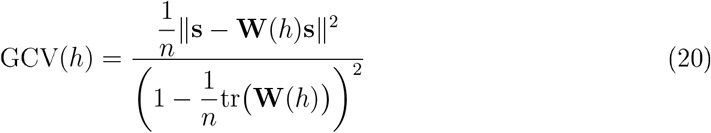

The numerator is the residual sum of squares (RSS) measuring the fit to the observed data, while the denominator penalizes model complexity through tr(**W**), the effective number of parameters. This formulation approximates the leave-one-out cross-validation error without the computational burden of *n*-fold resampling (Golub et al., 1979).

The optimal bandwidth is then:

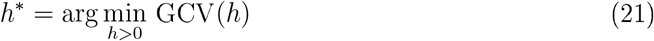

In practice, *h*^∗^ is found by evaluating GCV(*h*) over a logarithmically spaced grid of candidate bandwidths spanning the range from the median nearest-neighbor distance to the maximum spatial extent of the electrode array.

The NW quilting procedure consists of the following steps:

- **Aggregate sensor power:** For each subject *i* and each electrode *k*, compute the sensor-level power 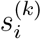 from the recorded time series (e.g., sum of squared voltages or spectral power).
- **Standardize coordinates:** Z-score all electrode positions 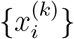 and cortical vertex coordinates {*v*} using the global electrode statistics.
- **Compute kernel weights:** For every (electrode, vertex) pair, evaluate the Gaussian kernel *K*_*h*_ using Equation (16).
- **Interpolate:** Apply the NW estimator (Equation 17) to obtain the estimated source power 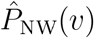 at each cortical vertex.

The NW estimator treats all sensor observations equally regardless of subject identity, effectively ignoring the hierarchical structure of the data. Consequently, it cannot disentangle inter-subject amplitude scaling differences from true spatial power variations. If subject *i* has a high overall gain factor, its electrodes will disproportionately in ate the estimated power at nearby vertices, producing a group map biased toward high-gain subjects. This is precisely the confound that the IOLMM framework resolves via explicit random-effects modeling of subject-specific scaling factors.

## Footnotes

1 Spurious source leakage derived from the ill-posed inverse operator acting on noise or distal signals.

